# Dynamics of phosphofructokinase condensation are regulated by metabolic and redox cues

**DOI:** 10.64898/2026.06.09.731068

**Authors:** Polina Reichert, Snezhana Oliferenko, Fabrice Caudron

**Affiliations:** School of Biological and Behavioural Sciences, Queen Mary University of London, Mile End Road, London E1 4NS, UK; The Francis Crick Institute, 1 Midland Road, London NW1 1AT, UK; Randall Centre for Cell and Molecular Biophysics, School of Basic and Medical Biosciences, King’s College London, Guy’s Campus, London, UK; School of Molecular Biosciences, University of Glasgow, G12 8QQ, Glasgow, UK; IGMM, Univ Montpellier, CNRS, 34293 Montpellier, France

## Abstract

Condensation of phosphofructokinase, the rate-limiting enzyme of glycolysis, into granules has been observed across species and in various stress conditions. However, it remains unclear what mechanisms govern this process. Here, we show that the two subunits of yeast phosphofructokinase, Pfk1 and Pfk2, assemble into granules even in the absence of exogenous stress, challenging the notion that PFK condensation in yeast is restricted to quiescent or stressed cells. While phosphofructokinase granules are predominant in replicatively aged cells, they are independent of cell division and can form before or after cells cease dividing. We further demonstrate that both subunits can co-localise to the same granules and that Pfk2 requires the disordered N-terminus of Pfk1 for assembly. Strikingly, phosphofructokinase granules are largely absent in cells engaged in respiration or lacking respiratory capacity, but their formation is strongly induced during the metabolic transition from fermentation to respiration. This suggests that phosphofructokinase condensation is tightly linked to the metabolic strategy of the cell. Moreover, we find that reactive oxygen species signaling, mediated by superoxide dismutase 1 (Sod1), modulates phosphofructokinase granule formation, as cells deficient in Sod1 exhibit impaired granule assembly. Overall, our results indicate that phosphofructokinase condensation is a dynamic process regulated by metabolic and redox cues.

## Introduction

The spatial regulation of metabolism has emerged as an important control of cellular physiology. In addition to localising to specific cellular organelles, enzymes from the same pathway may form membrane-less compartments (Bar-Peled and Kory, 2022; Chen and Silver, 2012; Gu et al., 2023). In the case of mammalian purinosomes that assemble under purine-depleted conditions, a colocalization of several enzymes controlling subsequent steps in purine biosynthesis increases metabolic output, replenishing purine levels (An et al., 2008). Other examples of enzyme condensation are assemblies of just one, critical enzyme within a pathway, such as phosphofructokinase (PFKL) filaments in rat mammary adenocarcinoma cells (Webb et al., 2017). In addition to their role as active sites of catalysis, enzyme assemblies have also been suggested to act as storage depots or facilitate degradation. In many cases their functions remain unknown. Remarkably, a screen in *Saccharomyces cerevisiae* has uncovered that more than 100 cytosolic proteins form reversible assemblies in response to changes in metabolite conditions (Narayanaswamy et al., 2009). Enzyme condensation thus appears to be a wide-spread and evolutionarily conserved mechanism for metabolic regulation.

As the gatekeeper of glycolysis, phosphofructokinase sets the pace of central carbon metabolism and is tightly controlled on multiple levels (Compton and Patrick, 2025; Hofmann, 1978; Yuan et al., 2025). Its catalytic function, the phosphorylation of fructose-6-phosphate, commits upstream sugar phosphates to glycolysis, resulting in the downstream production of ATP, NADH and crucial building blocks for the biosynthesis of cellular anabolites. Phosphofructokinase therefore controls the availability of pyruvate for fermentation in the cytosol or oxidative phosphorylation (OXPHOS) in mitochondria. As the interplay between the two major ATP generating pathways, glycolysis and OXPHOS, is tightly regulated, phosphofructokinase is subject to feedback inhibition by pyruvate and the TCA cycle intermediate citrate. Although phosphofructokinase is known to be regulated by over 20 metabolites, its spatial regulation remains poorly understood (Hicks et al., 2023; Lynch et al., 2024; Compton and Patrick, 2025).

The structure of phosphofructokinase is highly conserved throughout evolution. In yeast, phosphofructokinase hetero-octamers composed of the homologous Pfk1 and Pfk2 subunits ensure optimal function, but each subunit can also be active in the monomeric and other oligomeric forms (Compton and Patrick, 2025; Klinder et al., 1998). Both Pfk1 and Pfk2 have been shown to form filaments *in vivo* (Shen et al., 2016) and multiple studies showed their co-localisation with other glycolytic enzymes in the so-called glycolytic bodies (G-bodies). These dynamic assemblies accelerate the rate of glycolysis when OXPHOS function is compromised upon oxygen limitation (Jin et al., 2017; Miura et al., 2013). This phenomenon appears to be evolutionarily conserved as *C. elegans* neurons contain phosphofructokinase granules near synapses during transient anoxia and even in normoxic conditions during energy stress or following neuronal stimulation. Disrupting granule assembly impairs synaptic vesicle cycling, synaptic recovery and locomotion (Jang et al., 2016). Similarly, in *Drosophila* flight muscles, the co-localisation of glycolytic enzymes in sarcomeres is essential for flight, further highlighting the functional importance of enzyme condensation (Wojtas et al., 1997).

Despite these studies, how phosphofructokinase assemblies are formed and which cues trigger their formation in unstressed conditions remains unclear. To answer these questions, we investigated the prevalence of phosphofructokinase granules across different growth phases in *S. cerevisiae*. We show that phosphofructokinase granule assembly depends on disordered regions within the Pfk1 subunit and serves to sequester a fraction of the enzyme away from the diffuse pool, allowing little to no interchange between them. Strikingly, cells undergoing a shift from fermentation to respiration preferentially form phosphofructokinase granules, while cells incapable of reprogramming their metabolism display extremely low granule levels. Overall, our data indicate that phosphofructokinase granules are a cell-intrinsic response to changing redox conditions and metabolic strategies.

## Results

### Yeast phosphofructokinase granules form in normoxic conditions throughout all growth phases

To investigate if the formation of phosphofructokinase granules in normoxia is restricted to specific, stressful, conditions, e.g., the stationary phase, we analysed the localisation of endogenously tagged Pfk1-GFP and Pfk2-GFP throughout the growth cycle. Interestingly, both GFP-tagged phosphofructokinase isoforms formed single bright granules during exponential growth, albeit at different levels (Figure 1A-B). Pfk1-GFP could be found in granules earlier than its sister subunit, showing highest granule abundance during exponential growth, while Pfk2-GFP displayed highest granule levels in stationary phase. Applying a fluorescence threshold to both Pfk1-GFP and Pfk2-GFP images, which distinguishes all granules in the exponential growth phases, revealed that while Pfk2 granules displayed strong signal throughout all growth stages, Pfk1 granules became dimmer once cells reached stationary phase (Figure 1A).

**Figure 1.**
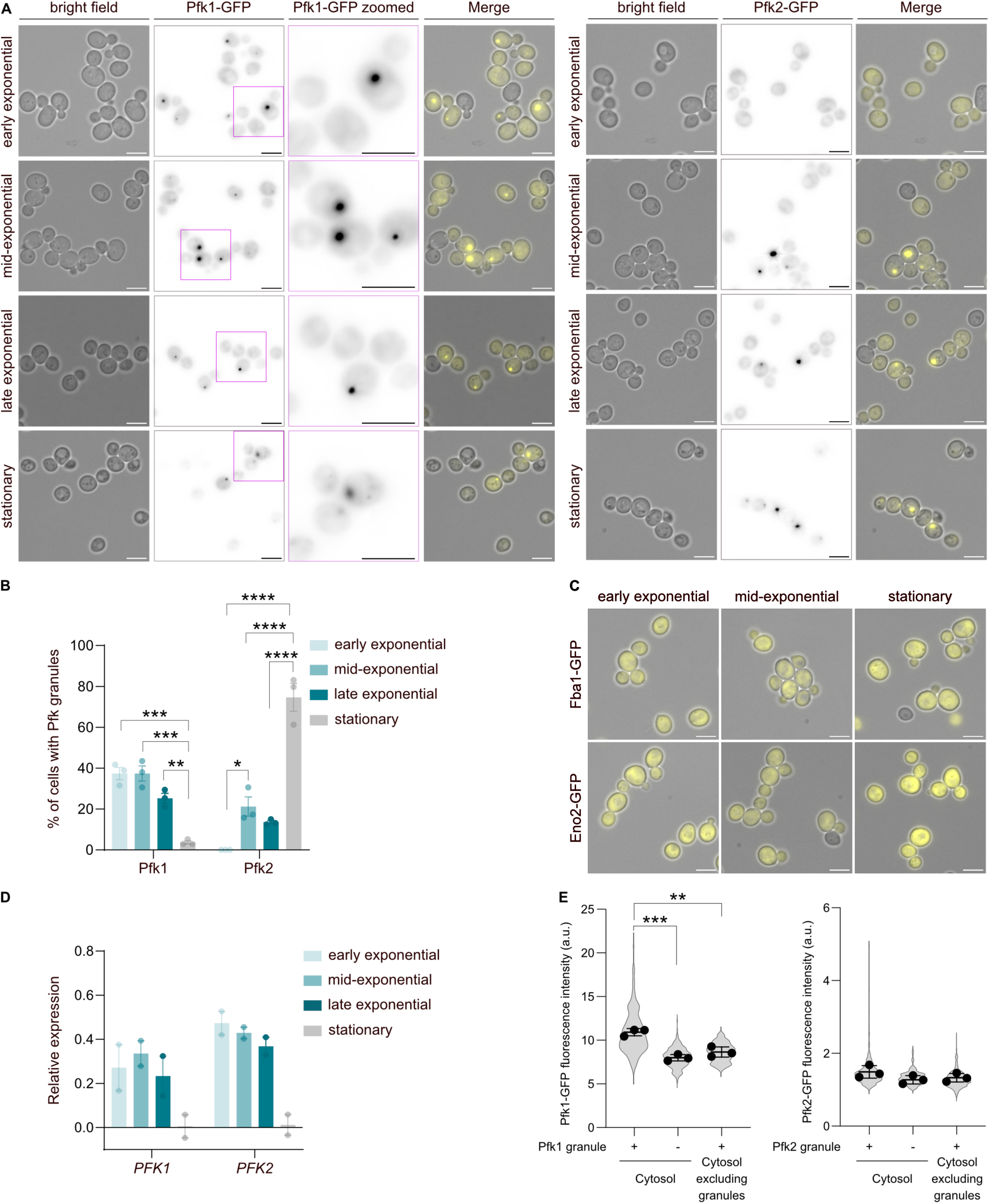
Pfk1 and Pfk2 show distinct granule profiles during normoxic growth. A) Localisation of Pfk1-GFP (left) and Pfk2-GFP (right) during exponential growth phases on glucose and in stationary cells. Scale bars = 5 µm. B) Percentage of cells with Pfk1-GFP and Pfk2-GFP granule levels across four growth stages (average ±SD). n > 100 cells per biological repeat, ***p<0.001, **** p < 0.0001 (One-way ANOVA). C) Diffuse distribution of aldolase (Fba1-GFP) and enolase (Eno2-GFP) in all growth stages. Scale bars = 5 µm. D) Pfk1 and Pfk2 transcript levels across exponential and stationary phases (average ±SD). E) Quantification of Pfk1-GFP and Pfk2-GFP fluorescence in cells with and without granules at exponential phase. The violin plots depicts the distributions of individual cells. The dots are the average of each replicate. Average and SD for the three replicates are shown. n > 100 cells per biological repeat, **p<0.01, ***p<0.001 (One-way ANOVA, n = 3 replicates).

Other glycolytic enzymes, such as aldolase (Fba1-GFP) and enolase (Eno2-GFP), displayed a diffuse distribution throughout all growth stages (Figure 1C). Transcript levels of both *PFK1* and *PFK2* remained stable during exponential growth but decreased in stationary phase, indicating that Pfk1 granule formation was underpinned by high gene expression levels (Figure 1D). Indeed, analysis of GFP fluorescence intensity in cells with and without granules at exponential phase shows that cells with Pfk1 granules display higher overall fluorescence, suggesting that Pfk1, but not Pfk2, granule occurrence coincides with higher protein levels (Figure 1E).

In our growth protocol, we used stringent culturing methods that leverage a longer recovery (over 12h) from dilution to minimise carry-over effects from the pre-culture and ensure that all cells are fully metabolically active. We then tested whether culturing methods that rely on imaging cells 2.5 hours (early exponential) and 5 hours (mid-exponential) after dilution from a saturated culture would influence granule levels. Both Pfk1 and Pfk2 could still be found in granules during exponential growth but the granule profile of Pfk1 now showed a strong spike in granule levels 5h after dilution (Figure S1A, B). Given that such a strong increase is not expected during exponential growth (Figure 1B), and we can exclude changes in gene expression as a driving factor (Figure S1C), we hypothesise that Pfk1 granule formation is concurrent with a shift in metabolism as cells fully recover from dilution shock and restart normal metabolic activity.

### The disordered N-termini of both subunits drive their localisation to phosphofructokinase granules

If the granules are important for metabolic reprogramming and serve as active sites of catalysis, we would expect them to contain both Pfk subunits to achieve highest Pfk activity. To reconcile the differing granule level profiles of Pfk1 and Pfk2, we next sought to determine whether they co-localise to the same granules. We therefore created a strain expressing Pfk1-mCherry and Pfk2-GFP from their endogenous loci and quantified their co-localisation in stationary phase, when Pfk2-GFP granules are abundant. The two subunits often accumulated in the same granules, with almost every Pfk1-mCherry granule showing Pfk2-GFP signal. Since we have previously observed that Pfk1-GFP granule fluorescence is decreased in stationary phase, it cannot be excluded that Pfk1 is present, albeit at much lower levels, in cells that appear to contain the Pfk2-GFP-’only’ granules (Figure 2A, B).

**Figure 2.**
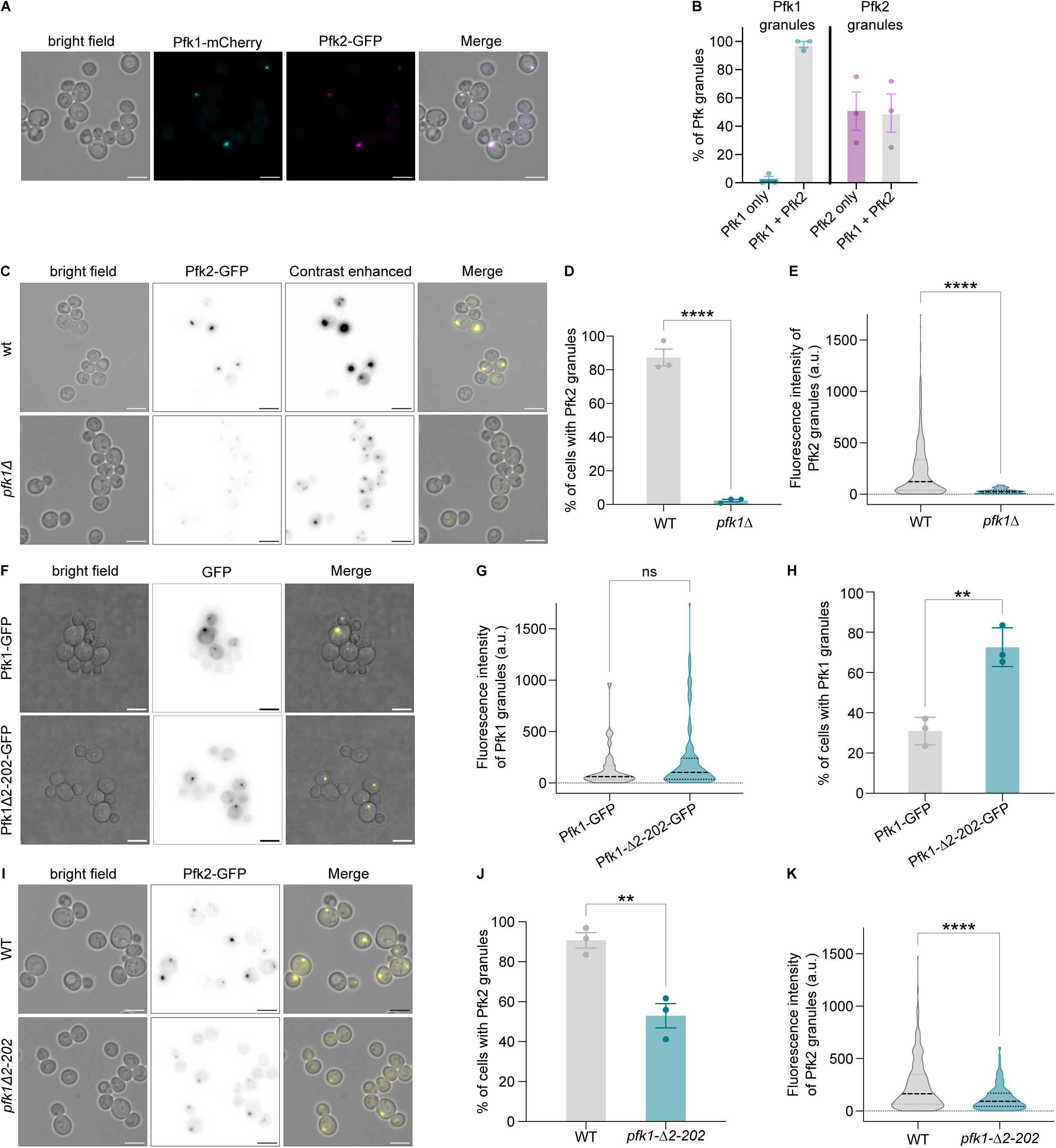
The recruitment of Pfk2 to Pfk1 granules is strongly influenced by the disorered N-terminus of Pfk1. A) Co-localisation of Pfk1-mCherry (cyan) and Pfk2-GFP (magenta) in granules in stationary phase. Scale bars = 5 µm. B) Left: quantification of Pfk1-mCherry granules that also co-localise or not with Pfk2-GFP granules. Right: quantification of Pfk2-GFP granules that also co-localise or not with Pfk1-mCherry granules (average ±SD). C) Localisation of Pfk2-GFP in wild type and *pfk1Δ* cells in stationary phase with normal (left) and enhanced (right) brightness. Scale bars = 5 µm. D) Percentage of cells with Pfk2-GFP granule in wild type and *pfk1Δ* cells in stationary phase (average ±SD). n > 100 cells per biological repeat, **** p < 0.0001 (t-test). E) Integrated intensity of Pfk2-GFP granules in wild type and *pfk1Δ* cells in stationary phase. n > 100 cells per biological repeat, ****p<0.0001 (Mann-Whitney test). F) Localisation of Pfk1-GFP and Pfk1-Δ2-202-GFP in stationary phase. Scale bars = 5 µm. G) Integrated intensity of Pfk1-GFP granules in *pfk1Δ* cells expressing full-length Pfk1-GFP or Pfk1Δ2-202-GFP in stationary phase. n > 100 cells per biological repeat. ns = not significant (t-test). H) Percentage of cells with Pfk1-GFP granule in wild type and *pfk1-Δ2-202* cells in stationary phase (average ±SD). n > 100 cells per biological repeat, ** p < 0.01 (t-test). I) Localisation of Pfk2-GFP in wild type and *pfk1-Δ2-202* cells in stationary phase. Scale bars = 5 µm. J) Percentage of cells with Pfk2 granules in *pfk1Δ* cells expressing full length Pfk1-GFP or Pfk1-Δ2-202-GFP cells in stationary phase (average ±SD). n > 100 cells per biological repeat, ** p < 0.01 (t-test). K) Integrated intensity of Pfk2-GFP granules in wild type and *pfk1Δ2-202* and *pfk1Δ* cells in stationary phase. n > 100 cells per biological repeat, **** p < 0.0001 (Mann-Whitney test).

Since our data suggest that Pfk1 is the first to condense into a granule that subsequently recruits Pfk2, we hypothesized that removing Pfk1 should impair the formation of Pfk2 granules. Indeed, knocking out *PFK1* led to a drastic reduction in the number of Pfk2-GFP granules that passed the fluorescence threshold (Figure 2C, D). Closer inspection revealed that Pfk2-GFP granules were present in most cells but were less intense (Figure 2E) and smaller (Figure S2A).

During hypoxia in yeast, the disordered N-terminus of Pfk2 is required for Pfk2 granule formation (Jin, M. et al., 2017). We tested whether a deletion of the disordered N-terminus of Pfk1 would have an impact on Pfk1 and Pfk2 granule formation. Cells expressing an N-terminally truncated form of Pfk1 (Pfk1-Δ2-202-GFP) were able to form Pfk1 granules of a similar intensity (Figure 2F-G) and size (Figure S2B) to cells expressing full-length Pfk1-GFP. Surprisingly, there were more cells with Pfk1-Δ2-202-GFP granules than with wild type Pfk1-GFP in stationary phase (Figure 2H), suggesting that Pfk1-Δ2-202-GFP granules are more stable than Pfk1-GFP granules. Therefore, Pfk1 does not fully require its disordered N-terminus to form granules. However, expression of *pfk1-Δ2-202* led to a decrease in the percentage of cells with Pfk2 granules (Figure 2I, J), as well as in their size (Figure S2C) and intensity (Figure 2K), yet this phenotype is not as strong as with a full knockout of *PFK1*. Overall, our data suggest that Pfk2, but not Pfk1, requires the disordered N-terminus of Pfk1 to condense into granules, and imply that Pfk1 may act as the granule nucleator.

### Dynamics of Pfk1 granules differ between subpopulations

We next sought to determine whether Pfk1 and Pfk2 exchange dynamically between the dilute cytosolic fraction and the condensed granule fraction. Fluorescence recovery after photobleaching (FRAP) experiments revealed two distinct behaviours within the pool of examined granules. About half of the granules (53%) showed no recruitment of unbleached protein into the partially-bleached granule, suggesting that they represent a pool of protein that does not easily exchange with the diffuse fraction. However, the other half of the examined granules recovered up to half their fluorescence within a 90-second time frame, indicating that these granules were capable of recruiting new protein (Figure 3A, B). The differences between the former, immobile and the latter, recovering fraction of granules might be due to differences in granule maturity and/or possibly function.

**Figure 3.**
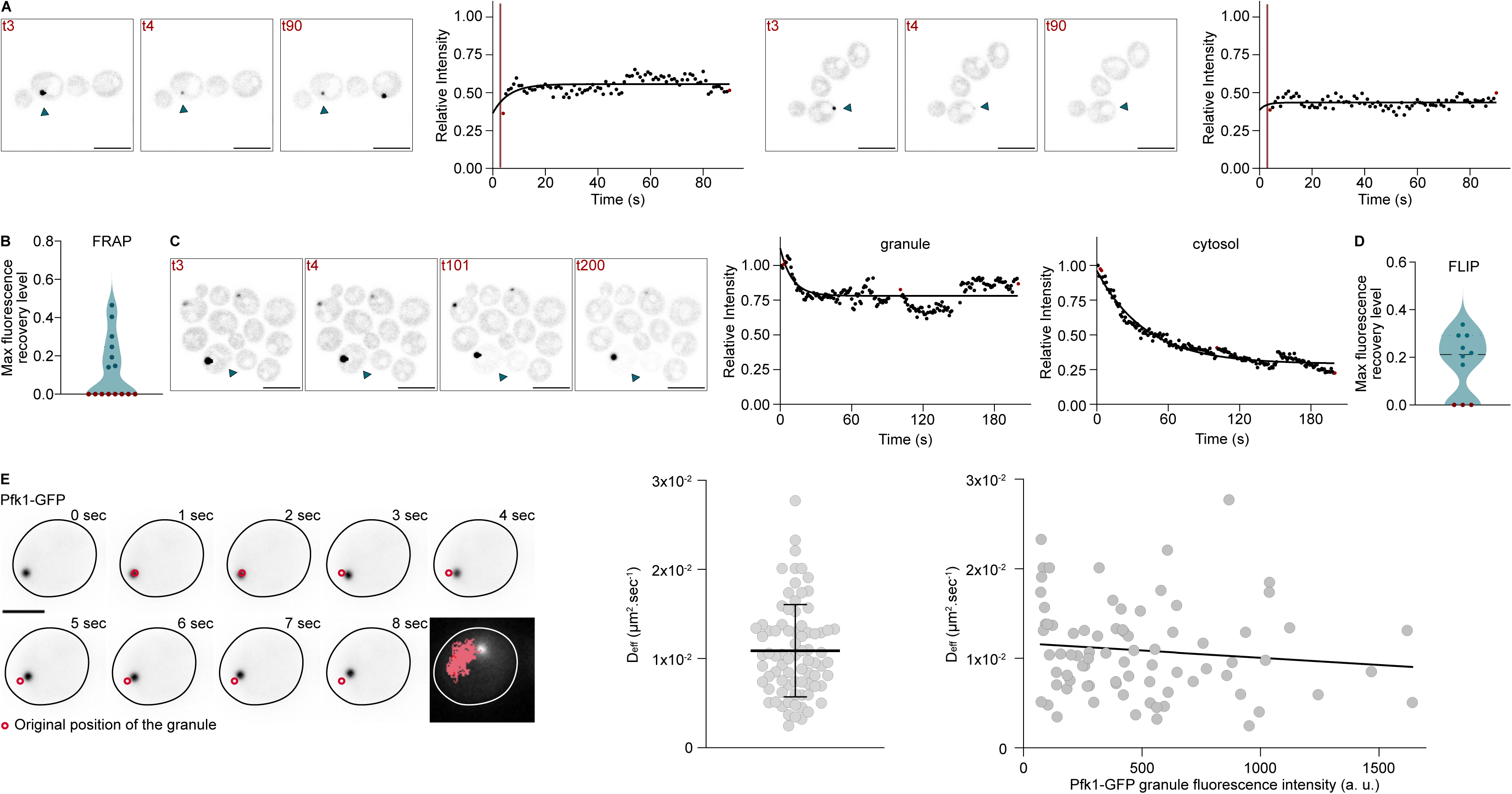
Dynamics of Pfk1 granules in glucose-grown cells. A) FRAP analysis after partial bleaching of Pfk1 granules at t = 3 seconds in cells indicated by arrowhead. Scale bars = 5 µm. Representative images of the two recovery profiles observed among cells across three time points (highlighted in red in the recovery graph on the right, n = 15 cells). The left cell is a representative profile of cells showing modest recovery. The right cell is a representative profile of cells showing no recovery. B) Quantification of maximum fluorescence recovery levels in FRAP experiment with the two subpopulations indicated in different colours. C) FLIP analysis following bleaching of the diffuse cytosolic signal in the cell indicated by arrowhead. Scale bars = 5 µm. Time points in fluorescence images indicated in red in relative fluorescence intensity profiles of the granule and cytosol (n = 10 cells). D) Quantification of maximum fluorescence loss levels in FLIP experiments with the two subpopulations indicated in different colours. E) Images of a movie used to track Pfk1-GFP granules. The original position of the granule at the beginning of the movie is displayed as a red circle, and the full track shown at the last image. Scale bar = 5 µm. The middle graph shows a scatter plot of individual effective diffusion for each granule tracked (n = 79 granules, average is shown ± SD). The right panel shows the distribution of the effective diffusion of individual granules as a function of the fluorescence intensity of the granules (A simple linear regression line is shown with an equation: Y = −1.613e-018*X + 1.168e-014).

Accordingly, fluorescence loss in photobleaching (FLIP) data showed two behaviours of protein exchange between granules and the diffuse pool. Upon bleaching of the cytosolic protein signal, the majority of granules (70%) displayed a decrease in fluorescence intensity. However, the highest observed loss accounted for only about a third of total granule intensity, suggesting that the exchange between the cytosolic and granule fractions is limited. The second fraction of granules showed no loss of fluorescence intensity, confirming that some Pfk granules do not exchange protein with the cytosolic pool (Figure 3C, D).

To determine whether Pfk1 granules were mobile and whether their size had any effect on their movement within the cell, we used widefield imaging to track granule diffusion. Neither the size nor the fluorescence intensity of Pfk1 granules appeared to have any influence on the effective diffusion of the granules (Figure 3E). Interestingly, Pfk1 granules displayed comparatively slow centre displacement, moving slower than would be expected of freely diffusing particles (previously shown to have diffusion coefficients of 0.2-0.4 (Delarue et al., 2018; Lemière and Chang, 2023)), which may be a reflection of their large size or suggest their attachment to a larger intracellular structure such as an organelle.

### Pfk1 granules are predominant in aged cells but are independent of cell division

We next asked if granules formed in a cell could be inherited by their daughter cells upon cell division. First, we tested whether Pfk1 granules persisted after inhibition of Hsp104, a chaperone essential for prion propagation. Hsp104 severs prion amyloid fibrils in mother cells, creating seeds that can diffuse to daughter cells, ensuring a stable inheritance of the prion state in the progeny (Byrne et al., 2007). Growing yeast cells in the presence of 3 mM guanidine HCl for multiple days to inhibit Hsp104 as previously described (Byrne et al., 2007) did not eliminate Pfk1 granules, showing that these granules do not behave as Hsp104-dependent prions (Figure S3).

Second, we attempted to observe events of granule inheritance by daughter cells using time-lapse microscopy. To facilitate the tracking of individual cells in a dividing population, we inhibited cell division in daughter cells using the Mother Enrichment Program (MEP) (Lindstrom and Gottschling, 2009). Addition of estradiol to the growth medium induces the knock-out of two essential genes (*UBC9* and *CDC20*), only in the daughter cells of the MEP strain. Therefore, upon estradiol addition, the mother cells continue to divide, while daughter cells permanently arrest in M-phase (Lindstrom and Gottschling, 2009). We tracked 70 dividing mother cells that exhibited Pfk1 granules over a total of 255 cell divisions and did not observe a single instance of granule inheritance (Figure 4A). We therefore conclude that granules are largely not inherited by daughter cells during cell division.

**Figure 4.**
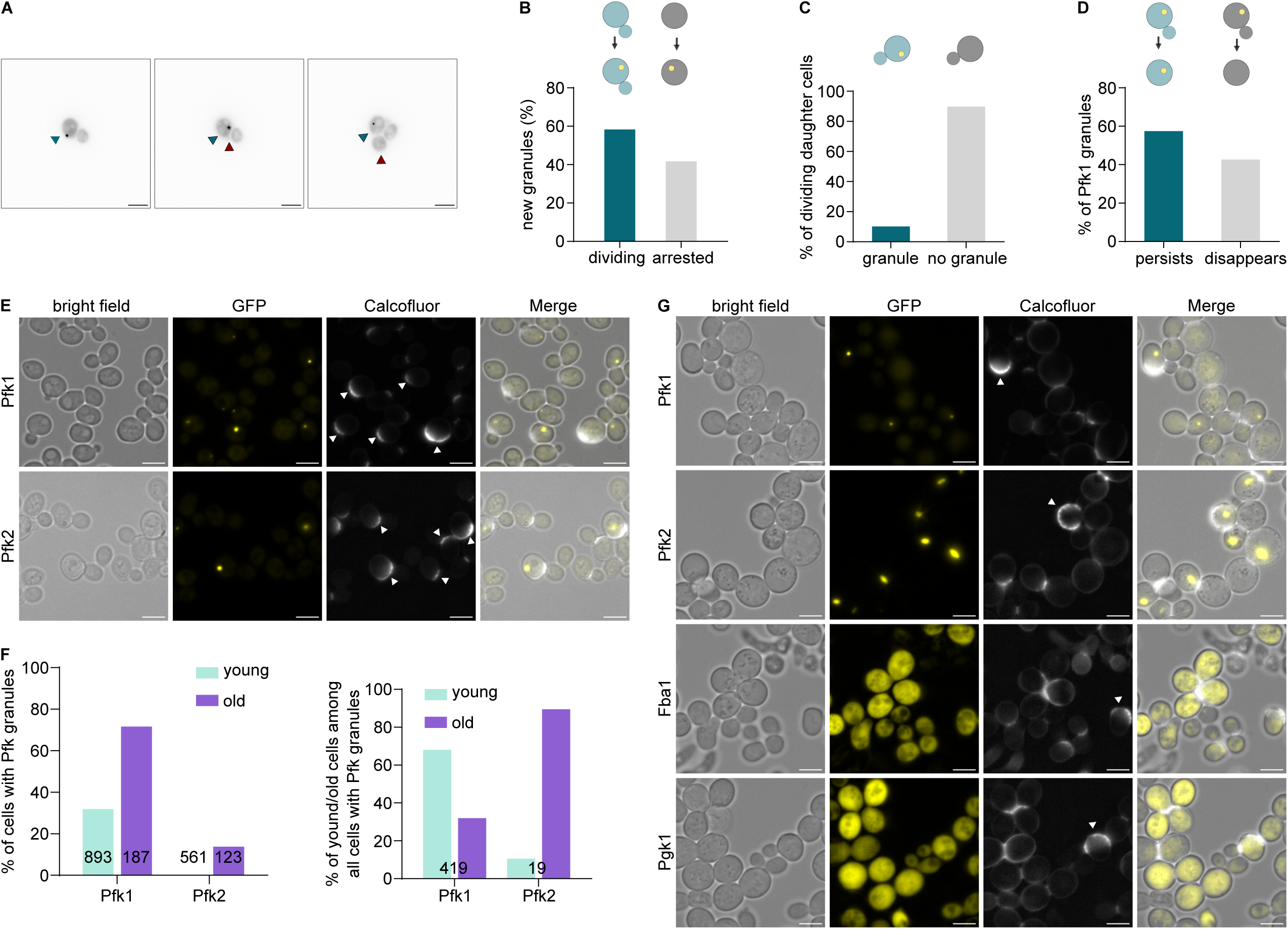
Phosphofructokinase granules are more common in replicatively aged cells but are not dependent on cell division. A) Time-lapse microscopy of a MEP cell dividing. Mother cell indicated by blue arrowhead, daughter cell indicated by red arrowhead. Scale bars = 5 µm. B) Quantification of new granules observed in dividing versus arrested MEP daughter cells using time-lapse microscopy. n = 70 cells. C) Quantification of dividing daughter cells with versus without granules using time-lapse microscopy. n = 70 cells. D) Quantification of the persistence of Pfk1 granules following the cessation of cell division using time-lapse microscopy. n = 70 cells. E) Calcofluor staining of exponentially growing Pfk1-GFP and Pfk2-GFP expressing cells. Old cells indicated by white arrowheads. Scale bars = 5 µm. F) Quantification of Pfk granules in young versus old cells shown in E. Left panel: Young versus old cells with granules out of all young/old cells. Right panel: Young versus old cells with granules out of all cells with granules. G) Calcofluor staining of MEP cultures of Pfk1-GFP, Pfk2-GFP, Fba1-GFP and Pgk1-GFP expressing cells. Old cells indicated by white arrowheads. Scale bars 5 µm.

As previous studies have shown that in normoxic conditions Pfk1 can be found in granules in stationary phase (Shen et al., 2016), we asked whether granule formation might be a phenomenon that is exclusive to cell cycle arrested cells. We leveraged the ability of some MEP daughter cells to escape cell cycle arrest for a few divisions to analyse whether new granules formed predominantly in dividing or truly arrested cells. The 48 new granules that formed within our imaging time frame were roughly evenly split between arrested and dividing daughter cells, showing that granules can form in both arrested and actively dividing cells (Figure 4B). Having established that cell division is not a prerequisite for granule formation, we flipped the question to ask whether granule formation might be a prerequisite for cell division. Within a pool of 274 daughter cells that divided only about 10% formed Pfk1 granules, indicating the lack of an interdependence between cell division and granule formation (Figure 4C).

Yeast mother cells can only produce a finite number of daughter cells until they enter senescence and ultimately die, a process known as replicative aging (Lin et al., 2025; Mortimer and Johnston, 1959). We used this phenomenon to analyse whether Pfk1 granules tend to persist past a mother cell’s last division and found that to be the case only in about half the cases, with the other half dissolving their granules before or around the time of the last cell division before arresting (Figure 4D). This result further suggests that granule formation is uncoupled from cell division.

To determine whether phosphofructokinase granules might be predominant among replicatively aged cells in an exponentially growing culture, we used calcofluor to visualise bud scars. Assessing the intensity of the calcofluor stained bud scars allowed us to gauge the stage within a cell’s replicative lifespan. For both Pfk1 and Pfk2, cells with granules represented a greater proportion within the aged cell population than among young cells (Figure 4E, F). As Pfk2 does not readily form granules during exponential growth (Figure 1), cells with granules are a relative rarity within both the aged and the young cell populations. Strikingly, within the population of all cells with Pfk2 granules, old cells by far outweighed young cells, representing 90% of all granule-carrying cells (Figure 4F). For Pfk1, the reverse is true: while 72% of all aged cells have Pfk1 granules (Figure 4E, F), aged cells with granules represent only 32% of the total pool of cells with granules (Figure 4F). Phosphofructokinase granules thus appear to be predominant in, but clearly not exclusive to, aged cells.

As MEP allows for the enrichment of aged cells in a cell population, we tested whether other glycolytic enzymes might form granules in aged cells as they have been shown to do in hypoxia (Jin et al., 2017; Miura et al., 2013). Tagging aldolase (Fba1) and phosphoglycerate kinase (Pgk1) with GFP in a MEP background strain did not show detectable granules in either arrested daughters or aged mother cells (Figure 4G). Granule formation under normoxia may therefore be a phenomenon that, within the glycolytic pathway, is exclusive to phosphofructokinase, independent of cell division but predominant in cells further along in their replicative lifespan.

### Pfk1 granule formation may be triggered by changes in the metabolic mode of a cell

Aged cells carry a higher oxidative burden (Brandes et al., 2013) - and given that glycolytic granules have previously been shown to form under hypoxia - we tested whether interfering with the cellular antioxidant defense system would have a discernible effect on Pfk1 granule formation. Jin and colleagues have shown that in hypoxic conditions glycolytic granule formation is completely abrogated in the absence of the Cu, Zn-superoxide dismutase 1 (Sod1) enzyme (Jin et al., 2017). Accordingly, we found that knocking out *SOD1* severely impaired granule formation in our experimental setup in normoxia (Figures 5A, S4A). To ensure that the observed phenotype was due to the lack of enzymatic activity of Sod1, we also analysed granule formation rates in the absence of the Sod1 copper chaperone Ccs1. We found that a *CCS1* knockout phenocopies the *SOD1* knockout and displays drastically reduced Pfk1 granule formation during exponential growth, suggesting that the cellular redox balance might be a key element in granule formation (Figures 5A, S4A).

**Figure 5.**
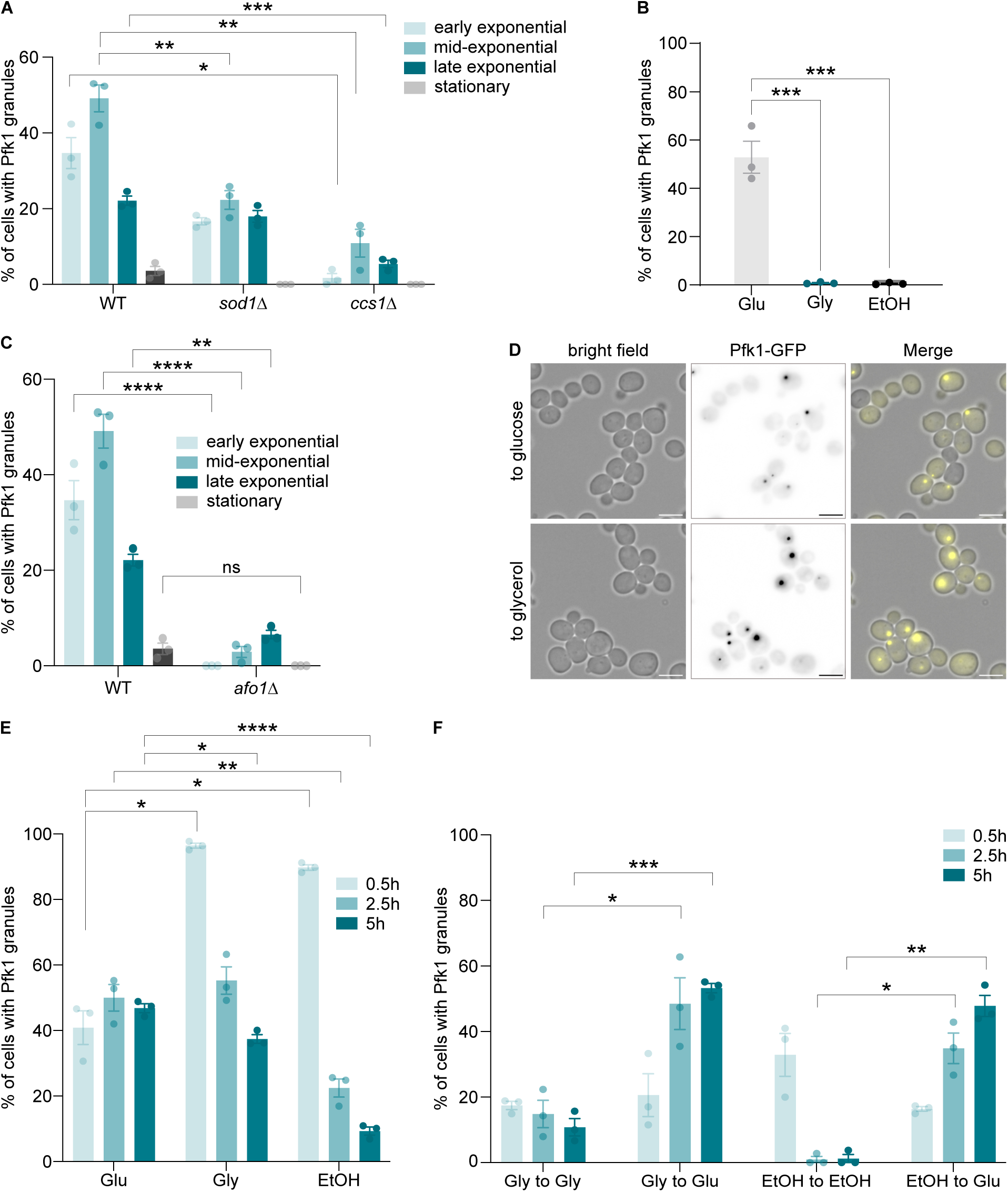
Phosphofructokinase granules depend on the metabolic mode of the cell. A) Percentage of wild type, *sod1Δ* and *ccs1Δ* cells with Pfk1-GFP granules across the growth curve (average ±SD). n > 100 cells per biological repeat, * p < 0.05, ** p < 0.01, *** p < 0.001 (One-way ANOVA). B) Percentage of cells grown on glucose, glycerol and ethanol with a Pfk1-GFP granule (average ±SD). n > 100 cells per biological repeat, *** p < 0.001 (One-way ANOVA). C) Percentage of wild type and *afo1Δ* cells with a Pfk1-GFP granule across the growth curve (average ±SD). n > 100 cells per biological repeat, ns = not significant, ** p < 0.01, **** p < 0.0001 (One-way ANOVA). D) Localisation of Pfk1-GFP into granules following carbon source substitution. Scale bars = 5 µm. E) Percentage of cells with Pfk1 granules following carbon source substitution (average ±SD). n > 100 cells per biological repeat, **** p < 0.0001 (One-way ANOVA). F) Percentage of cells with Pfk1 granules in cells following carbon source substitution from respiratory carbon sources to glucose (average ±SD). n > 100 cells per biological repeat, * p < 0.05, ** p < 0.01, *** p < 0.001, **** p < 0.0001 (One-way ANOVA).

Next, we sought to test the effect of higher reactive oxygen species (ROS) levels on the prevalence of phosphofructokinase granules. To do so, we grew cells in respiratory media (glycerol and ethanol) as respiratory metabolism is directly linked to an increase in mitochondrial ROS production (Andreyev et al., 2005; Murphy, 2009). Interestingly, we found that the incidence rate of Pfk1 granules in respiring cells (grown on glycerol or ethanol) was extremely low, even lower than in *sod1Δ* and *ccs1Δ* strains grown on glucose. This suggests that the metabolic strategy of a cell may be a key determinant in Pfk1 coalescence (Figures 5B, S4B).

To further test this hypothesis, we abrogated respiratory capacity in our yeast strain by knocking out Ageing factor 1 (Afo1), a mitochondrial ribosomal protein of the large subunit. *afo1Δ* strains have previously been shown to be respiratory deficient as they lack mitochondrial DNA and are incapable of growing on respiratory substrates. We chose to use the *afo1Δ* background instead of the commonly used rho^0^ background as *afo1Δ* cells, in contrast to rho^0^ cells, do not display a growth defect on fermentative substrates and have a cell size that is similar to wild type cells (Heeren et al., 2009). If metabolic mode alone were the decisive factor in granule formation, we would expect the *AFO1* knockout, which ensures that all cells are fermenting, to produce the opposite phenotype to that observed in respiring cells. Intriguingly, we found that both respiratory deficient and respiring cells have a drastically decreased incidence rate of Pfk1 granule formation (Figures 5C, S4C). The low Pfk1 granule levels in *afo1Δ* cells hint for the first time at an involvement of mitochondrial function, possibly beyond respiration *per se*, in the regulation of glycolytic enzyme coalescence.

Given that shifting the metabolic mode to either full respiration or full fermentation negatively impacted granule formation, we hypothesised that Pfk1 granule formation may be a response to changes in the intracellular environment that occur when cells reprogram their metabolic strategy. To test this hypothesis, we switched cells from growth on glucose to growth on respiratory substrates (glycerol and ethanol) and monitored their granule levels as they switched from a fermentative to a respiratory strategy. Strikingly, for both respiratory substrates, we saw a drastic upregulation of granule levels within 30 minutes of carbon source substitution that was not present in the glucose-to-glucose control condition. The spike in granule formation abated over the next 5 hours as cells fully switched to respiration, confirming that the low incidence rate of Pfk1 granules on respiratory media is indeed a phenotype of fully respiring cells (Figure 5D, E). As Pfk1 granule formation was not triggered when we forced cells to switch to fermentation by substituting respiratory substrates with glucose (Figure 5F), it appears that Pfk1 granules form in response to changes that occur when cells initiate respiration which may be yet another indication that the cellular redox balance is a driving factor in granule formation.

## Discussion

In line with previous studies, we found that yeast phosphofructokinase is capable of forming granules *in vivo*. Our findings show that its condensation into granules is not exclusive to stationary cultures or quiescent cells and can readily occur under normoxic conditions in the absence of exogenous stressors. These granules appear to contain both phosphofructokinase subunits, albeit at different levels given the dissimilar granule profiles of Pfk1 and Pfk2 in different growth phases (Figure 1). Some granules may only contain one of the two subunits or one of the subunits may be much less enriched in granules. As both subunits tend to maintain some level of diffuse distribution even when a large proportion of the protein localises to granules, the differing granule profiles likely reflect changing proportions of the two subunits within phosphofructokinase granules.

This is further supported by the findings that Pfk2 appears to depend on the presence of Pfk1 and especially its disordered N-terminus for proper granule formation. In the presence of an N-terminally truncated Pfk1 (*pfk1-Δ2-202*), Pfk2 shows an impairment in granule formation, characterised by fewer, dimmer and often smaller granules (Figure 2). As this is highly reminiscent of the phenotype observed under hypoxia when the N-terminus of Pfk2 itself was similarly truncated (Jin et al., 2017), it might hint at common biophysical interactions between the N-termini of the two subunits in both normoxic and hypoxic conditions. Interestingly, Pfk1 and Pfk2 fold and assemble into heterodimers co-translationally in budding yeast. While Pfk1 does not interact with Pfk2 until quite late in its translational process, Pfk2 associates with Pfk1 just after its disordered N-terminus emerges from the ribosome (Shiber et al., 2018), suggesting that the nascent N-terminus of Pfk1 might contain an assembly signal that is important for dimerization of the two subunits. Should this indeed be the case, it further implies that dimerization of the two subunits is imperative for Pfk2 to assemble into granules but, given our findings that the same truncation has no effect on Pfk1 localisation, it appears to be dispensable for Pfk1 condensation into granules (Figure 2). As the interaction between four Pfk1 subunits forms the core of the octameric form of phosphofructokinase (Banaszak et al., 2011), it may be that Pfk1 is prone to oligomerization, in a manner that is independent of its N-terminus, and that co-translational engagement with Pfk2 might prevent uncontrolled Pfk1 multimerization.

Taking into account our findings that a) Pfk1 granules are predominant in aged cells, b) interfering with the cellular redox balance decreases granule levels and c) respiratory deficient cells have lower granule levels, it would be tempting to speculate that an obvious cue for Pfk1 granule formation is a change in intracellular ROS production (Figures 4, 5). If granule formation depends on a cell’s ability to quickly sense changes in the local redox balance, both respiring cells and *sod1/ccs1*-deficient cells would be at a disadvantage as they routinely deal with higher ROS levels than redox-balanced, fermenting cells. Given that mitochondria heavily contribute to both the production and the detoxification of ROS, it is thus not surprising that *afo1Δ* cells which lack functional mitochondria have drastically reduced granule levels (Figure 5).

This ties in nicely with the observed spikes in granule formation following an initiation of respiration: the increase in mitochondrial ROS as a by-product of respiration would then be responsible for the increase in Pfk1 granules. The predominance of granules in aged cells could potentially be explained in the same way: aged cells have recently been suggested to adopt a respiratory metabolism, regardless of the carbon source in the media (Leupold et al., 2019), so will have undergone a switch from fermentation to respiration. The decrease in Pfk1 granules in the first few hours after the initial spike following the initiation of respiration in young cells may be due to yet again changing intracellular redox conditions as cells mount the antioxidant defence. Aged cells, on the other hand, have a reduced capacity to defend against ROS, which might explain the persistence of granules (Figure 5E).

Hypoxia might be yet another condition of local, intracellular changes in the redox balance as hypoxia-induced ROS production has been widely reported in a range of organisms including yeast (Dirmeier et al., 2002; Guzy et al., 2007). It may be that the phosphofructokinase granules reported here, the glycolytic granules (‘G-bodies’) in hypoxic yeast and possibly even the phosphofructokinase granules observed in *C. elegans* neurons during transient anoxia, share common mechanisms of formation. The first report of the formation of glycolytic granules in yeast showed that interfering with the redox balance in hypoxic conditions, by treatment with either a reductant (N-acetylcysteine) or ROS inducers (antimycin A or CCCP), impairs granule formation, further cementing the importance of ROS homeostasis in the induction of granules (Miura et al., 2013). While it may be that phosphofructokinase senses changes in the local redox balance indirectly via ROS signaling, it is also possible that the cue for granule assembly is direct enzyme oxidation, particularly of the single cysteines found in the N-termini of both subunits. Using ROS to affect the main rate-setting enzyme of glycolysis would be an elegant way of mediating metabolic signalling between mitochondria and the cytosol.

The apparent absence of other glycolytic enzymes in the phosphofructokinase granules described here presents a major difference to the glycolytic granules observed in previous studies in hypoxic conditions. Given the circumstances surrounding their formation, it is possible that the granules reported here and glycolytic granules in hypoxia also differ in their function. In hypoxia, glycolysis becomes the main strategy for ATP production and multiple previous studies have reported both that other ATP consuming enzymes co-localise to these granules and that these granules appear to be important for higher glycolytic output (Jin et al., 2017; Miura et al., 2013). As our data suggest that Pfk2 depends on the presence of Pfk1 to properly form granules, it implies a level of co-regulation of the two subunits in phosphofructokinase granules in normoxia.

Conversely, the apparent absence of other glycolytic enzymes in the granules, their induction during a switch to respiration and their limited numbers in highly fermenting respiratory-deficient cells might hint at other functions of phosphofructokinase granules in normoxia. They may serve as storage depots that ensure that a fraction of protein is protected from degradation in case respiration does not initiate successfully or they may provide a means of spatially segregating lower glycolysis from other pathways that branch off its upstream metabolites. For example, it could serve to release phosphofructokinase’s control of glycolysis globally within the cell, allowing more glucose-6-phosphase to enter the oxidative branch of the pentose phosphate pathway and increasing NADPH levels to combat oncoming oxidative stress. In cells that are switching to respiration, they could furthermore serve to facilitate the switch from glycolysis to gluconeogenesis by spatially sequestering the enzyme responsible for one of the main irreversible glycolytic reactions. Whatever the precise function of the phosphofructokinase granules described here is, a sequestration of a large proportion of the gatekeeper of glycolysis (accounting for a similar or even higher amount of protein than is distributed diffusely in the cytosol (Figure 1E)) is bound to affect which metabolic pathways predominate in the cell.

Another interesting question about phosphofructokinase granules in normoxia that deserves further attention is whether they are RNA-dependent. Indeed, the phosphofructokinase β-subunit Pfk2 was recently shown to possess RNA unwinding activity and to modulate cell cycle progression (Albihlal et al., 2026), supporting a direct role for RNA in granule assembly and suggesting potential connections between metabolic regulation and cell division. Hypoxic G-bodies have been shown to contain RNA, and specifically mRNAs encoding glycolytic enzymes (Fuller et al., 2020; Jin et al., 2017). The first study investigating glycolytic foci in hypoxia has found their formation to be inhibited upon treatment with the translational inhibitor cycloheximide, suggesting an active role of mRNA translation in their assembly (Miura et al., 2013). An imaging approach has furthermore revealed the presence of core-fermentation (‘CoFe’) granules in actively fermenting cells that are composed of glycolytic mRNAs undergoing active translation (Morales-Polanco et al., 2021). As most glycolytic enzymes have moonlighting functions as RNA-binding proteins and can bind their own mRNAs (Castello et al., 2015), it would be fascinating to investigate whether a cessation in active glycolytic mRNA translation upon induction of respiration, aids in the assembly of phosphofructokinase granules. Given the apparent importance of the disordered N-termini of the two subunits for granule formation and the possibility of their involvement in the co-translational enzyme oligomerization described above, probing the role of phosphofructokinase translation in granule assembly is a promising future direction. Alternatively, it would be interesting to probe whether metabolic enzyme assemblies, in a fashion reminiscent of stress granules, serve to store metabolic proteins and mRNAs to protect them against irreversible oxidative damage and subsequent degradation.

Overall, it is clear that phosphofructokinase granules are far more prevalent during normal growth in normoxic conditions than previously observed. Our work highlights the importance of carefully selecting culturing conditions to ensure that all cells are fully metabolically active, as we obtained drastically different granule profiles when giving cells less time to recover from dilution shock (Figure S1). Using our growth protocols, we show that these granules appear to be seeded by the Pfk1 subunit, are independent of cell division and occur more readily in aged cells. As they are induced when cells switch their metabolic program from fermentation to respiration and are strongly downregulated in both respiratory-deficient and oxidatively stressed cells, we suggest that a change in the intracellular redox balance may serve as a cue for granule formation. The high proportion of protein sequestered into such granules implies that they may have a strong effect on the efficiency of the glycolytic pathway and future work will have to elucidate the function of these granules in normoxic conditions.

## Materials & Methods

### Yeast husbandry and molecular genetics

All strains used in this study were derived from s288c BY4741 wild type (yFC01: *MAT***a**, *his3*Δ*1, leu2*Δ*0, ura3*Δ*0, met15*Δ*0*). The Mother Enrichment Program background (Lindstrom and Gottschling, 2009) was used to obtain aged cells. Gene deletions and C-terminal fusions were performed according to Janke et al., 2004. N-terminal deletions were obtained using the *URA3* pop-in/pop-out and counter-selection system (Schneider et al., 1995). All strains used in this study are listed in Table 1.

**Table 1.** Strains used in this study.

| Strain | Strain | Genotype |
| --- | --- | --- |
| PR1 | wild type | *his3Δ1 leu2Δ0 ura3Δ0 met15Δ0 MATa* |
| PR2 | Pfk1-GFP | *his3Δ1 leu2Δ0 ura3Δ0 met15Δ0 MATa PFK1-GFP:HIS3* |
| PR3 | Pfk2-GFP | *his3Δ1 leu2Δ0 ura3Δ0 met15Δ0 MATa PFK2-GFP:HIS3* |
| PR4 | Fba1-GFP | *his3Δ1 leu2Δ0 ura3Δ0 met15Δ0 MATa FBA1-GFP:HIS3* |
| PR5 | Eno2-GFP | *his3Δ1 leu2Δ0 ura3Δ0 met15Δ0 MATa ENO2-GFP:HIS3* |
| PR6 | Pfk1-mCherry,<br>Pfk2-GFP | *his3Δ1 leu2Δ0 ura3Δ0 met15Δ0 MATa*<br>*PFK1-mCHERRY:kanMX PFK2-GFP:HIS3* |
| PR7 | *pfk1Δ* Pfk2-GFP | *his3Δ1 leu2Δ0 ura3Δ0 met15Δ0 MATa pfk1::kanMX*<br>*PFK2-GFP:HIS3* |
| PR8 | *pfk1Δ1-202*<br>Pfk2-GFP | *his3Δ1 leu2Δ0 ura3Δ0 met15Δ0 MATa pfk1-Δ1-202:3HA*<br>*PFK2-GFP:HIS3* |
| PR9 | MEP Pfk1-GFP | *ade2::hisG his3 leu2 lys2 ura3D0 trp1D63 MATa*<br>*hoD::SCW11prCre-EBD78-NatMX loxP-UBC9-loxP-LEU2*<br>*loxP-CDC20-IntronloxP-HPHMX PFK1-GFP:HIS3* |
| PR10 | MEP Pfk2-GFP | *ade2::hisG his3 leu2 lys2 ura3D0 trp1D63 MATa*<br>*hoD::SCW11prCre-EBD78-NatMX loxP-UBC9-loxP-LEU2*<br>*loxP-CDC20-IntronloxP-HPHMX PFK2-GFP:HIS3* |
| PR11 | MEP Fba1-GFP | *ade2::hisG his3 leu2 lys2 ura3D0 trp1D63 MATa*<br>*hoD::SCW11prCre-EBD78-NatMX loxP-UBC9-loxP-LEU2*<br>*loxP-CDC20-IntronloxP-HPHMX FBA1-GFP:HIS3* |
| PR12 | MEP Pgk1-GFP | *ade2::hisG his3 leu2 lys2 ura3D0 trp1D63 MATa*<br>*hoD::SCW11prCre-EBD78-NatMX loxP-UBC9-loxP-LEU2*<br>*loxP-CDC20-IntronloxP-HPHMX PGK1-GFP:HIS3* |
| PR13 | *afo1Δ* Pfk1-GFP | *his3Δ1 leu2Δ0 ura3Δ0 met15Δ0 MATa afo1::kanMX*<br>*PFK1-GFP:HIS3* |
| PR14 | *sod1Δ* Pfk1-GFP | *his3Δ1 leu2Δ0 ura3Δ0 met15Δ0 MATa sod1::kanMX* |
|  |  | *PFK1-GFP:HIS3* |
| PR15 | *ccs1Δ Pfk1-GFP* | *his3Δ1 leu2Δ0 ura3Δ0 met15Δ0 MATa ccs1::kanMX*<br>*PFK1-GFP:HIS3* |

**Table 2.**
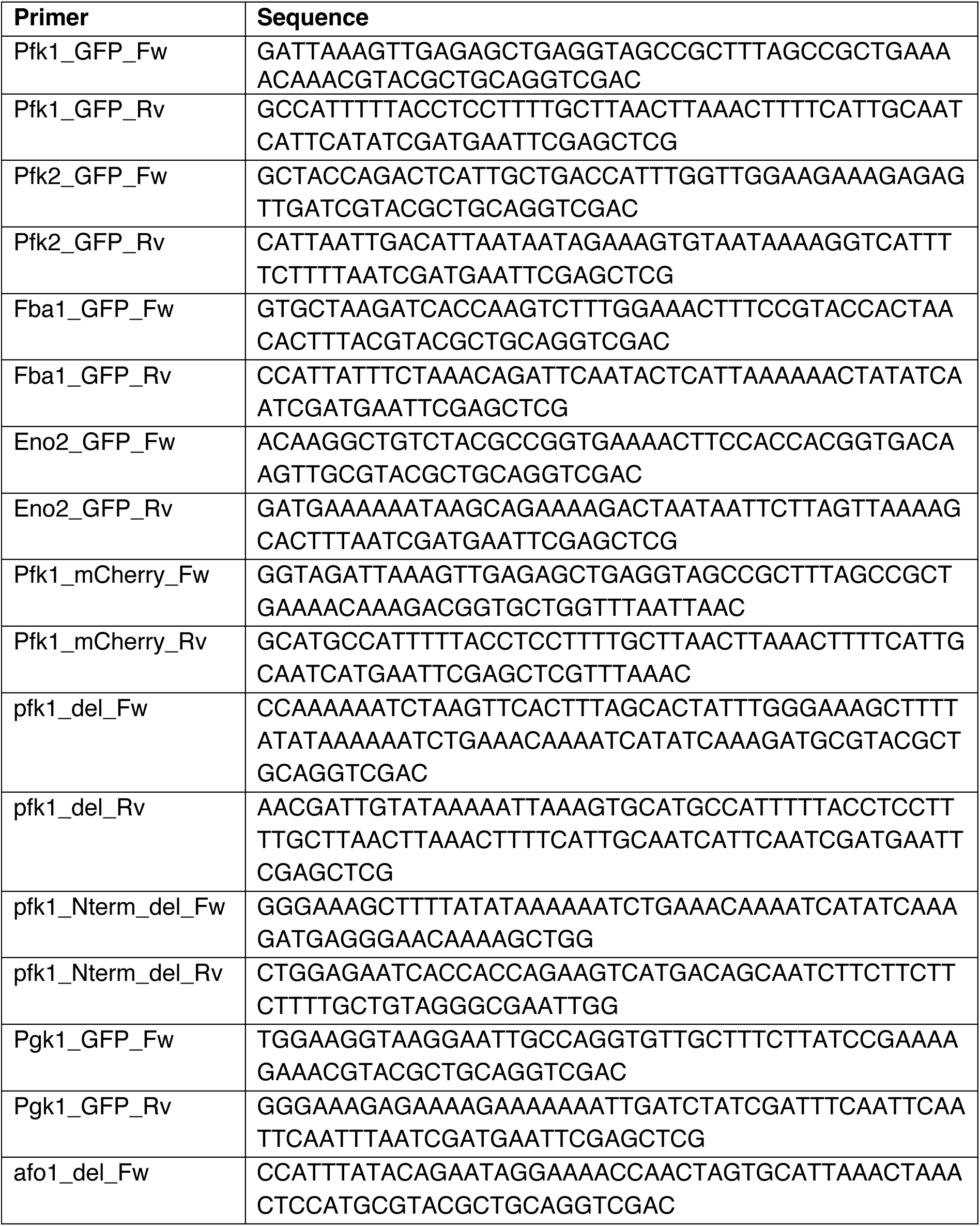
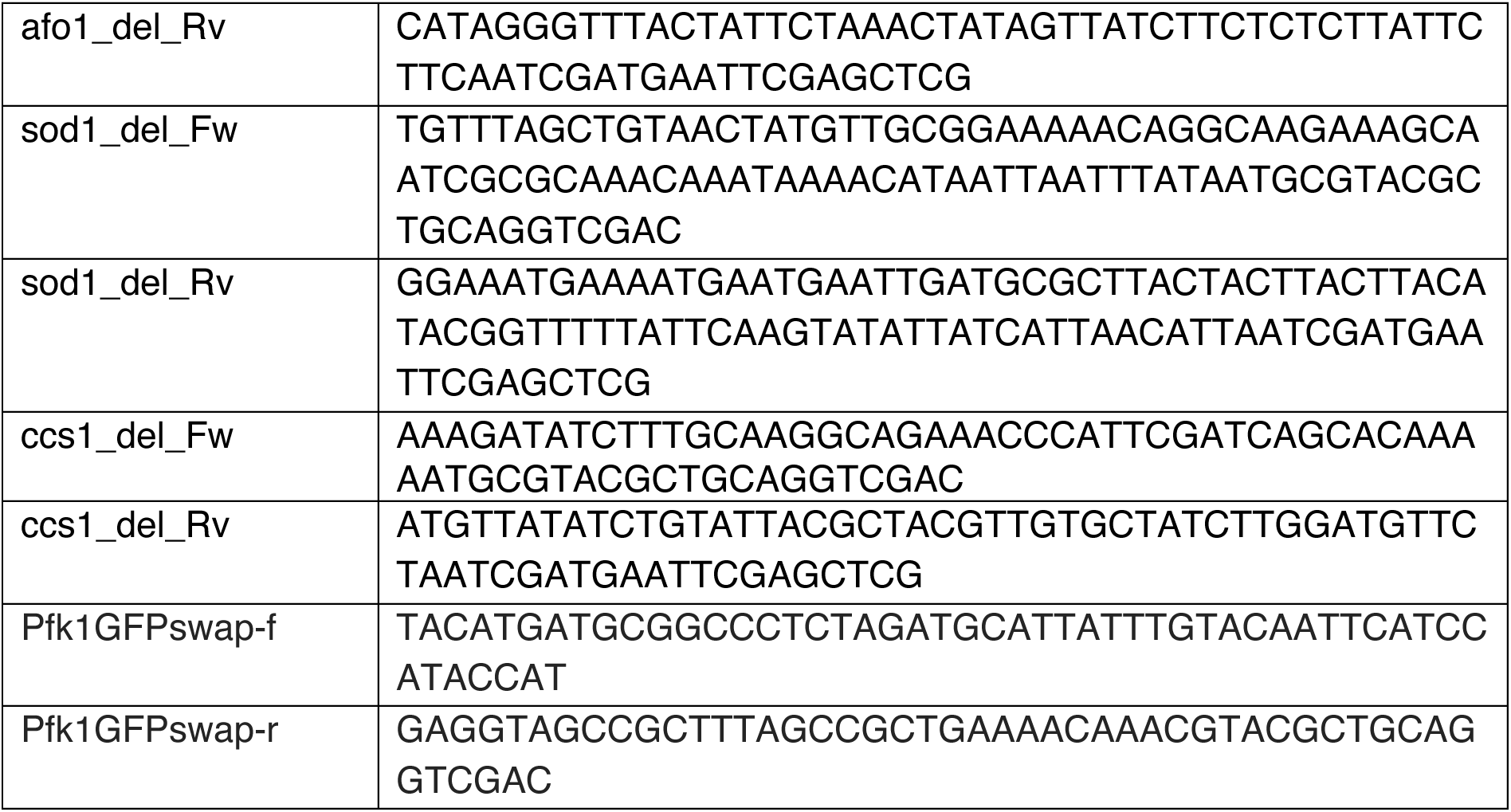
Primers used in this study.

### Growth and treatment of yeast

Yeast cells were grown in yeast extract peptone dextrose (YPD, 1% (w/v) yeast extract, 2% (w/v) peptone, 2% (w/v) glucose, 0.04% (w/v) adenine) or synthetic complete medium (SC, 6.7 g/L yeast nitrogen base, 0.77 g/L synthetic complete mix, 0.04% (w/v) adenine, supplemented with 2% (w/v) glucose (glucose medium) or 3% glycerol and 0.1% (w/v) glucose or 3% ethanol and 0.1% (w/v) glucose (ethanol medium). Calcofluor (FB28) staining was conducted by addition of the dye to 1 mL of cells at OD_595_ = 0.8 in YPD to a final concentration of 3 µg/mL and incubation for 5 min in the dark. Cells were washed three times and imaged immediately. For guanidine HCl treatment, cells were re-streaked on YPD agar containing 3mM guanidine HCl for four consecutive days before being imaged.

To study granule prevalence in aged cells, the Mother Enrichment Program was induced at an OD_595_ = 0.02 by the addition of 1 µM β-estradiol. Cells were grown for 17h, harvested by centrifugation and imaged. For time lapse microscopy of aged cells, cells were imaged every 20min on SCD agar containing 1 µM β-estradiol for 40h.

### Confocal microscopy

Cells were imaged on agar pads composed of SCD medium using a Zeiss Axio Observer Z1 microscope fitted with an α Plan-FLUAR 100x/1.45 NA objective lens and Hamamatsu Orca-Flash4.0 C11440 camera. Images are max projections of Z-stacks with background subtraction. For granule quantification, images were similarly thresholded.

Time lapse microscopy was performed using a Deltavision Elite (GE Healthcare) microscope equipped with a sCMOS camera and solid-state light-emitting diodes controlled by Softworx. A fluorescein isothiocyanate filter was used for imaging GFP fluorescence. Z-stack resolved deconvolution was performed using Softworx. Images were acquired every 20 min over a time span of 40h.

FRAP and FLIP experiments were performed using a Zeiss LSM880 NLO, imaging cells every 3 seconds, with an Apochromat 63x/1.4 NA Oil DIC M27 immersion objective. Fluorophores were bleached using the 488 nm line of the Argon laser.

For FRAP experiments images were acquired every second for 90 seconds, with 3 consecutive bleaches performed after the first 3 seconds of recording. Fluorescence intensity of the bleached region of interest (FRAP ROI) and a reference region in an adjacent cell was quantified and background signal (quantified in an area with no cells) was subtracted. Fluorescence intensity in the FRAP ROI was normalised to the reference region using the following formula (Bolognesi et al., 2016):

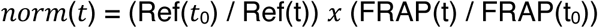

Ref(t_0_) = fluorescence intensity in reference region before bleaching
Ref(t) = fluorescence intensity in reference region at time point t
FRAP(t) = fluorescence intensity in FRAP ROI at time point t
FRAP(t_0_) = fluorescence intensity in FRAP ROI before bleaching

For FLIP experiments images were acquired every second for 200sec with continuous bleaching. Fluorescence of the granule (granule ROI) in the bleached cell was quantified, as was fluorescence of the diffuse signal in the bleached cell (diffuse ROI). Reference regions were taken from three cells in the same field of view. Background was subtracted from all quantified fluorescence intensities. The reference regions were averaged and the fluorescence intensity of the granule and the diffuse ROI was normalised to the bleaching of the reference regions. The fluorescence intensity profiles were plotted using one-phase decay with least squares fit (constraint: K > 0).

### Widefield microscopy

Time-lapse experiments of PFK1/2 were acquired on a Nikon Ti2 wide-field microscope, using a Nikon Plan Apo 100X/1.4 NA objective, every 1000 ms with a Photometrics Prime sCMOS camera.

### Cloning

Centromeric plasmids expressing *PFK1* or *pfk1-ΔN202* fused to mCherry under the control of the *PFK1* promoter and followed by the *CYC1* terminator were purchased from VectorBuilder. Both vectors were cut with XhoI and BamHI to remove mCherry. GFP was amplified from pFC22 (pYM44) using primers Pfk1GFPswap-f and Pfk1GFPswap-r. The PCR product was inserted into the cut vectors by homologous recombination using the NEBuilder HiFi DNA Assembly kit (New England Biolabs) to swap mCherry with GFP. Vectors were sequenced to validate the constructs.

### Real time quantitative PCR (qPCR)

RNA extraction was performed using the RNAeasy Mini Kit (Qiagen). cDNA was synthesized from 500ng of total RNA using the Transcriptor First Strand cDNA synthesis kit (Roche Diagnostics) and random hexamer primers. The qPCRBIO SyGreen Blue Mix (PCR Biosystems) was used on 25ng of cDNA and the reactions were run on a QuantStudio 6 instrument (Applied Biosystems). Cycle conditions used for amplification were as follows: 95°C for 2 min, followed by 40 cycles consisting of 5 sec at 95°C and 20 sec at 60°C. Using the Comparative C_T_ (ΔΔC_T_) method, the data was normalised to stationary phase with UBC6 as the reference gene.

## Acknowledgements

We acknowledge the imaging facility MRI, member of the national infrastructure France-BioImaging (https://ror.org/01y7vt929) supported by the French National Research Agency (ANR-24-INBS-0005 FBI BIOGEN) and the “Yeast media and technologies service” of IGMM and CRBM for providing us with ready-to-use media. We would like to thank Rocco D’Antuono and the Light Microscopy STP at the Francis Crick Institute for help with confocal and wide field microscopy. P.R was supported through a PhD fellowship from London Interdisciplinary Doctoral Programme (LIDo). F.C. was supported by MUSE funding. This work was supported by the Francis Crick Institute, which receives its core funding from Cancer Research UK (CC0102), the UK Medical Research Council (CC0102), and the Wellcome Trust (CC0102), and the Wellcome Trust Senior Investigator Award (103741/Z/14/Z) and Wellcome Trust Investigator Award in Science (220790/Z/20/Z) to S. O.

---

**Figure S1.**
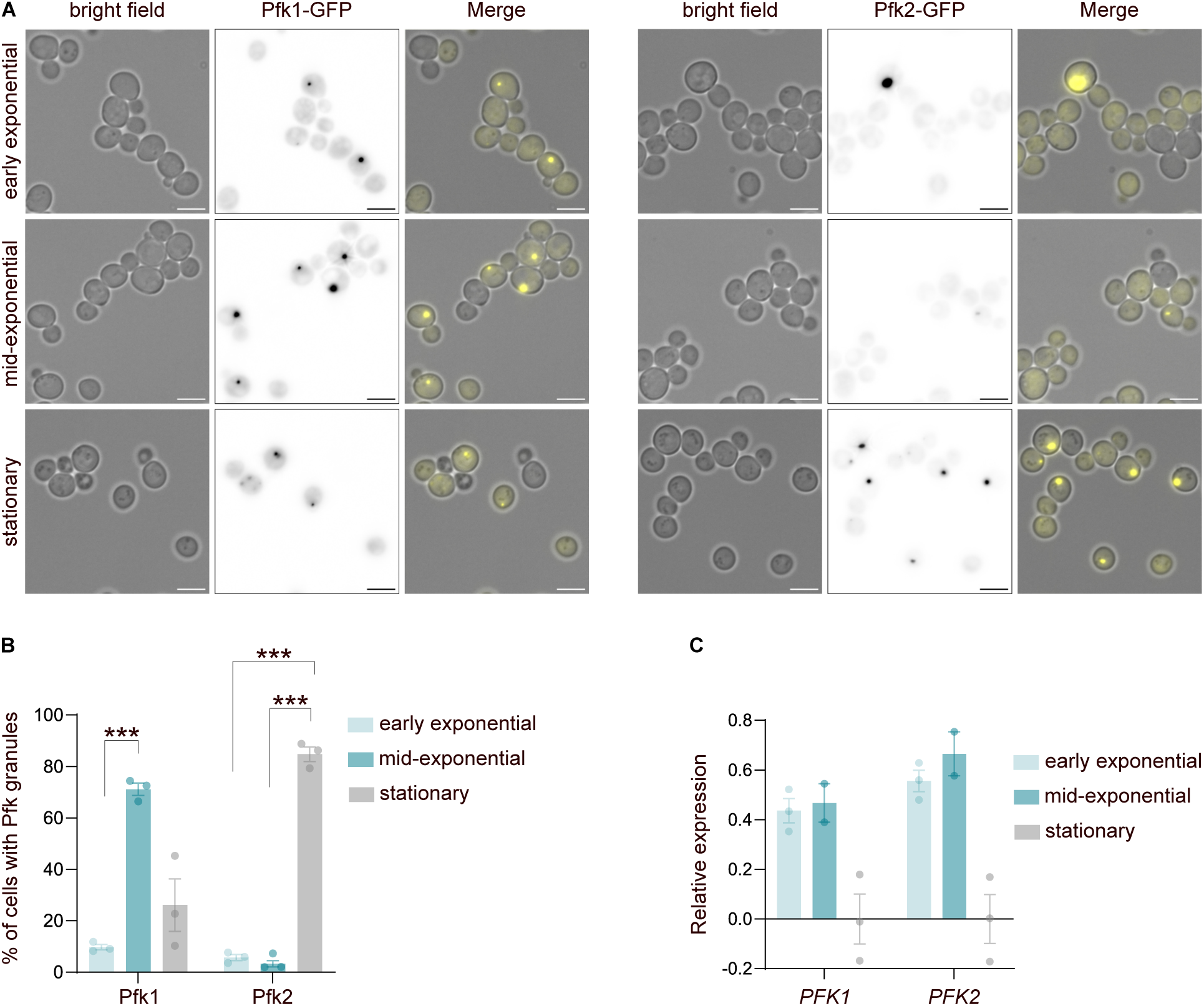
Effect of different culturing methods on phosphofructokinase coalescence. A) Localisation of Pfk1-GFP and Pfk2-GFP granules 2.5h (early exponential phase) and 5h (mid-exponential phase) after dilution and in stationary phase. Scale bars 5 µm. B) Percentage of cells with a Pfk1-GFP or Pfk2-GFP granule across growth stages (average ±SD). n > 100 cells per biological repeat, ***p<0.001 (One-way ANOVA). C) Pfk1 and Pfk2 transcript levels across exponential and stationary phases (average ±SD).

**Figure S2.**
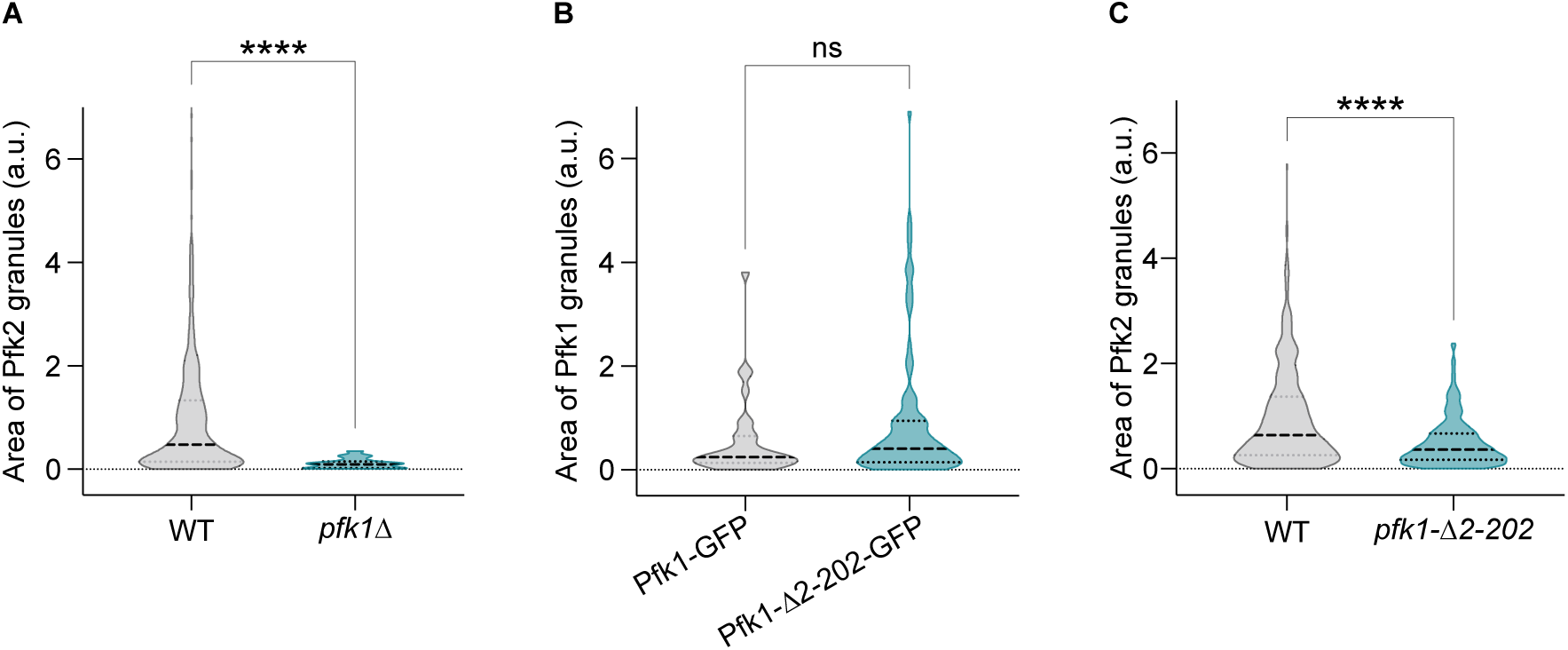
Area of phosphofructokinase granules in wild type and *PFK1* mutants. A) Area of Pfk2-GFP granules in wild type and *pfk1Δ* cells in stationary phase. n > 100 cells per biological repeat, ****p<0.0001 (Mann-Whitney test). B) Area of Pfk1-GFP granules in *pfk1Δ* cells expressing full-length Pfk1-GFP or Pfk1Δ2-202-GFP in stationary phase. ns = not significant (t-test). C) Area of Pfk2-GFP granules in wild type and *pfk1Δ2-202* and *pfk1Δ* cells in stationary phase. n > 100 cells per biological repeat, **** p < 0.0001 (Mann-Whitney test).

**Figure S3.**
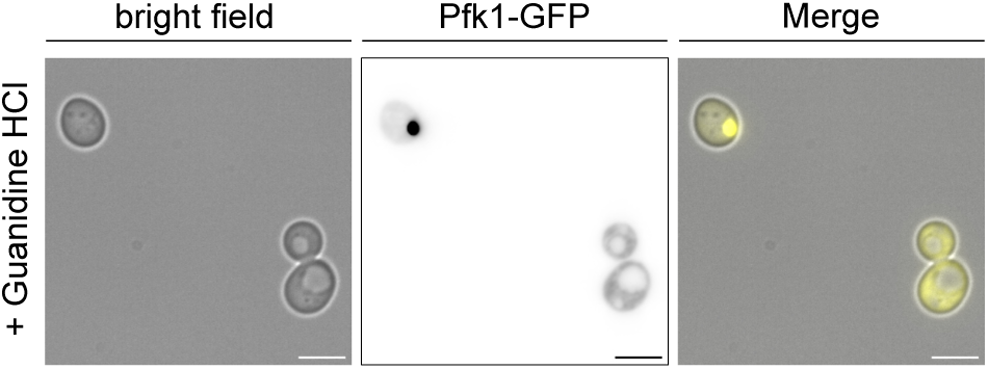
Hsp104 inhibition has no effect on Pfk1 granules. Localisation of Pfk1-GFP into granules after treatment with 3 mM guanidine HCl. Scale bars = 5 µm.

**Figure S4.**
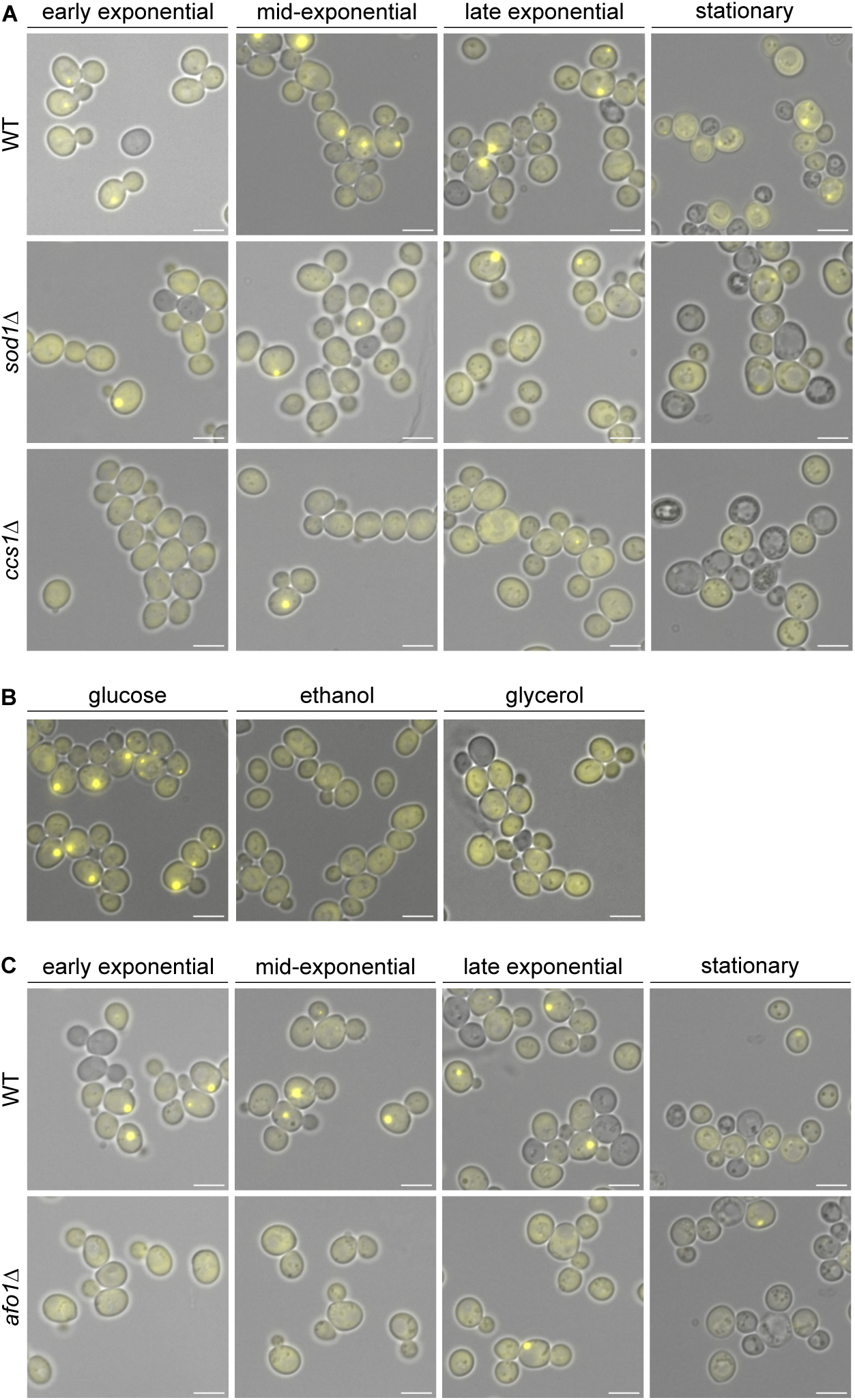
Localisation of Pfk1-GFP in Sod1 deficient mutant cells, respiratory conditions and respiratory mutant cells. A) Localisation of Pfk1-GFP during exponential growth and stationary phases in WT, *sod1Δ* and *ccs1Δ* mutant cells. Scale bars = 5 µm. B) Localisation of Pfk1-GFP in stationary phase in cells grown in fermentative (glucose) and respiratory (ethanol and glycerol) media. Scale bars = 5 µm. C) Localisation of Pfk1-GFP during exponential growth and stationary phase in WT and *afo1Δ* cells. Scale bars = 5 µm.

